# A combined bioinformatics and LC-MS based approach for the development and benchmarking of a comprehensive database for CNS proteins in *Lymnaea stagnalis*

**DOI:** 10.1101/2021.05.03.442491

**Authors:** Sarah Wooller, Aikaterini Anagnostopoulou, Benno Kuropka, Michael Crossley, Paul R. Benjamin, Frances Pearl, Ildikó Kemenes, György Kemenes, Murat Eravci

**Affiliations:** Bioinformatics Group, School of Life Sciences, University of Sussex, Brighton, United Kingdom; Sussex Neuroscience, School of Life Sciences, University of Sussex, Brighton, United Kingdom; Institute for Chemistry and Biochemistry, Freie Universität Berlin, Berlin, Germany

**Keywords:** *Lymnaea* Central Nervous System, Liquid Chromatography–Mass Spectrometry, bioinformatics, new proteomics database

## Abstract

Applications of key technologies in biomedical research, such as qRT-PCR or LC-MS based proteomics, are generating large biological (-omics) data sets which are useful for the identification and quantification of biomarkers involved in molecular mechanisms of any research area of interest. Genome, transcriptome and proteome databases are already available for a number of model organisms including vertebrates and invertebrates. However, there is insufficient information available for protein sequences of certain invertebrates, such as the great pond snail *Lymnaea stagnalis*, a model organism that has been used highly successfully in elucidating evolutionarily conserved mechanisms of learning and memory, ageing and age-related as well as amyloid-β induced memory decline. In this investigation, we used a bioinformatics approach to designing and benchmarking a comprehensive CNS proteomics database (LymCNS-PDB) for the identification of proteins from the Central Nervous System (CNS) of *Lymnaea stagnalis* by LC-MS based proteomics. LymCNS-PDB was created by using the Trinity TransDecoder bioinformatics tool to translate amino acid sequences from mRNA transcript assemblies obtained from an existing published *Lymnaea stagnalis* transcriptomics database. The blast-style MMSeq2 software was used to match all translated sequences to sequences for molluscan proteins (including *Lymnaea stagnalis* and other molluscs) available from UniProtKB. LymCNS-PDB, which contains 9,628 identified matched proteins, was then benchmarked by performing LC-MS based proteomics analysis with proteins isolated from the CNS of *Lymnaea stagnalis*. MS/MS analysis using the LymCNS-PDB database led to the identification of 3,810 proteins while only 982 proteins were identified by using a non-specific Molluscan database. LymCNS-PDB provides a valuable tool that will enable us to perform quantitative proteomics analysis to identify a plethora of protein interactomes involved in several CNS functions in *Lymnaea stagnalis* including learning and memory, aging-related memory decline and others.

## Introduction

Protein networks are performing key functions in all living organisms. The need for studying such complex biological functions has led to the development of LC-MS based platforms that are enabling researchers to perform quantitative system-wide analysis of proteomes, including protein-protein interactions, post translational modifications and spatial localization of proteins, even at the single cell level. Furthermore, several labelling and pre-fractionation techniques enable researchers to improve the detection of low abundance and possibly functional relevant proteins.

The great pond snail, *Lymnaea stagnalis*, is used as a model organism in a wide range of biological research fields, such as the study of host–parasite interactions, ecotoxicology, evolution, developmental biology, learning and memory, ageing and age-related memory decline, genome editing, ‘omics’ and human disease modelling (Benjamin et al., 2021; Benjamin and Kemenes, 2020; Fodor et al., 2020; Rivi et al., 2020).

In our laboratory, we are interested in elucidating evolutionarily conserved molecular mechanisms involved in the formation of long-term memory after classical conditioning. We already have established important roles for a variety of key enzyme proteins, such as NOS (Kemenes et al., 2002), MAPK (Ribeiro et al., 2005), PKA (Kemenes et al., 2006) and CaMKII (Naskar et al., 2014) and the transcription factors CREB1 (Ribeiro et al., 2003) and C/EBP (Hatakeyama et al., 2006). However, the lack of comprehensive proteomics information in *Lymnaea* has been hindering further progress with research aimed at understanding the roles of these proteins in the context of large-scale protein networks in the central nervous system.

Although, qRT-PCR is a well-established method to quantify specific gene transcripts of interest, it does not allow the quantification of a functional protein in its protein network, or arrive at conclusions about its activity due to possible post-translational modifications, e.g., phosphorylation.

To enable us to provide these types of important information, in the present study we have performed large-scale proteomics experiments by using nanoLC-MS to analyse protein expression and post translational modifications in the CNS of *Lymnaea stagnalis*.

One important prerequisite for a successful proteomic workflow and the accurate identification of proteins is the existence of a protein sequence database of the organism of interest that can be used for the comparison of the peptide sequences, acquired from the MS/MS fragmentation spectra with the protein sequences in the database of the same organism.

In UniProtKB (The UniProt Consortium 2019; (UniProt, 2019)), the only available and useful protein database for *Lymnaea stagnalis* consists of 519 proteins, of which 48 are already reviewed but 471 are still un-reviewed (Uniprot.org - Last modified on August 3, 2020).

Because of this lack of specific protein sequence information, previous proteomics analysis of *Lymnaea stagnalis* were performed by utilising the entire UniprotKB/Swissprot database with protein sequences from all available organisms (Rosenegger et al., 2010; Silverman-Gavrila et al., 2011) or using a Metazoa specific database (Giusti et al., 2013) to identify proteins.

To prepare a more comprehensive and representative database for CNS proteins of *Lymnaea stagnalis*, which will enable us to accurately identify proteins by LC-MS/MS analysis, we have selected the transcriptome dataset of Sadamoto *et al*., 2012 that is among the available transcriptome datasets (Bouetard et al., 2012; Davison and Blaxter, 2005; Dong et al., 2021; Feng et al., 2009; Sadamoto et al., 2012) published from the NCBI database and contains the transcriptome of the whole CNS including the buccal “learning” ganglia and the central cerebral ring.

The transcripts from the Sadamoto et al. 2012 dataset (NCBI accession number PRJDB98) were filtered for coding regions and the remaining transcripts were then searched for homology against all UniProtKB molluscan entries.

The resulting protein database, LymCNS-PDB, was then utilized in a proof of principle experiment by performing LC-MS based proteomics analysis with proteins isolated from the CNS of *Lymnaea stagnalis*. Furthermore, we have prepared a DB with all proteins from the LymCNS-PDB using the matching amino acid sequences of the other molluscan species from UniProtKB. With both databases, having the same size, we were able to compare the number of identifications from a *Lymnaea stagnalis* specific database and with a non-specific molluscan database.

## Materials and Methods

### Experimental animals

Specimens of *Lymnaea stagnalis* were raised in the breeding facility of the University of Sussex, where they were kept in 20–22°C copper free water under 12 h light and dark cycle. They were fed on lettuce 3 times and a vegetable-based fish food twice a week.

### Preparation of snail CNS

Whole CNS of 90 snails at the age of three months weighing approx. 1.5 g were prepared as follows: The shell of the snail was cut, and the animal’s body carefully removed. The dissected preparation was then pinned down in a Sylgard-coated dish containing HEPES buffered saline and dissected under a stereomicroscope (E-Zoom6, Edmund Optics, Barrington NJ, USA). The CNS was accessed by an incision in the dorsal body region isolated from the buccal mass by the severing of all the peripheral nerves, and then immediately placed in Eppendorf tubes on dry ice. Three tubes, with each containing 30 CNS, were stored at -80°C.

### Protein extraction, digestion and prefractionation

Frozen CNS samples were thawed for 30 min at room temperature (20–22°C) before adding lysis buffer (6M urea / 2M thiourea in 10mM HEPES pH 8.0) and 30 ceramic beads (1.4 mm zirconium oxide beads) for homogenisation in a Precellys 24 Homogeniser (Bertin Instruments) using two cycles for 30 seconds at 6800 RPM with a 60 second break in between, followed by a centrifugation for 1h at 14000 x g to remove the debris. The supernatant was transferred to a fresh tube and the extracted proteins were reduced for 30 min in 10 mM dithiothreitol and alkylated for 30 min in 55 mM iodoacetamide (in the dark). Proteins were first pre-digested with LysC for at least 3h at RT and after dilution with 3 volumes of 50 mM ammonium bicarbonate buffer, the main digestion was performed with trypsin overnight at RT. Peptide samples (triplicates) were desalted with C18 Hypersep cartridges (Thermo Fisher) and eluates were concentrated in a SpeedVac concentrator (Savant) and prefractionated into 6 fractions using the IPG strip-based peptide fractionation method as previously described by (Eravci et al., 2014).

### LC-MS analysis

Triplicates of 6 desalted peptide fractions (18 samples in total) were analysed by a reversed-phase capillary nano liquid chromatography system (Ultimate 3000, Thermo Scientific) connected to a Q Exactive HF mass spectrometer (Thermo Scientific). Samples were injected and concentrated on a trap column (PepMap100 C18, 3 µm, 100 Å, 75 µm i.d. x 2 cm, Thermo Scientific) equilibrated with 0.05 % trifluoroacetic acid in water. After switching the trap column inline, LC separations were performed on a capillary column (Acclaim PepMap100 C18, 2 µm, 100 Å, 75 µm i.d. x 25 cm, Thermo Scientific) at an eluent flow rate of 300 nl/min. Mobile phase A contained 0.1 % formic acid in water, and mobile phase B contained 0.1 % formic acid in 80 % acetonitrile, 20 % water. The column was pre-equilibrated with 5 % mobile phase B and peptides were separated using a gradient of 5–44% mobile phase B within 100 min. Mass spectra were acquired in a data-dependent mode utilising a single MS survey scan (m/z 350–1650) with a resolution of 60,000 in the Orbitrap, and MS/MS scans of the 15 most intense precursor ions with a resolution of 15,000. HCD-fragmentation was performed for all ions with charge states of 2+ to 5+ normalized collision energy of 27 and an isolation window of 1.4 m/z. The dynamic exclusion time was set to 20 s. Automatic gain control (AGC) was set to 3×106 for MS scans using a maximum injection time of 20 ms. For MS2 scans the AGC target was set to 1×105 with a maximum injection time of 25 ms.

MS and MS/MS raw data were analysed using the MaxQuant software package (version 1.6.12.0) with an implemented Andromeda peptide search engine (Tyanova et al., 2016). Data were searched against the FASTA formatted protein database of *Lymnaea stagnalis* described in the present manuscript.

### Database construction

To predict coding regions in the NCBI PRJDB98 transcriptome dataset, we analysed all available transcript assemblies with Trinity TransDecoder (v 5.5.0).

By then we had downloaded all available swissprot and trembl amino acid sequences for molluscan proteins from UniProtKB and characterized them by whether they were sequences for *Lymnaea stagnalis*, other euthyneura species, or other molluscan species.

All sequences were compared with the transdecoded database from the NCBI PRJDB98 transcriptome dataset mentioned above, in order to find potential matches using the blast-style program MMSeqs2 (Steinegger and Soding, 2017).

In each case, proteins were matched by preference, first against *Lymnaea stagnalis*, then other euthyneura and lastly against other molluscan proteins.

Duplicate Proteins as well as matches to proteins termed ‘uncharacterised’ or ‘hypothetical’ were removed from the final data set.

For all matching entries the header of the trembl/swissprot protein was used to update the headings of the respective uncharacterised *Lymnaea stagnalis* amino acid sequences from the transdecoded PRJDB98 transcriptome dataset, indicating the original GenBank accession number, the UniProt ID and the protein name of the matching molluscan protein and the p-value for matching accuracy provided by the MMSeqs2.

The constructed database LymCNS-PDB is saved in a ‘FASTA’ format following the UniProtKB parsing rules for FASTA headers (see Figure 1).

**Figure 1.**
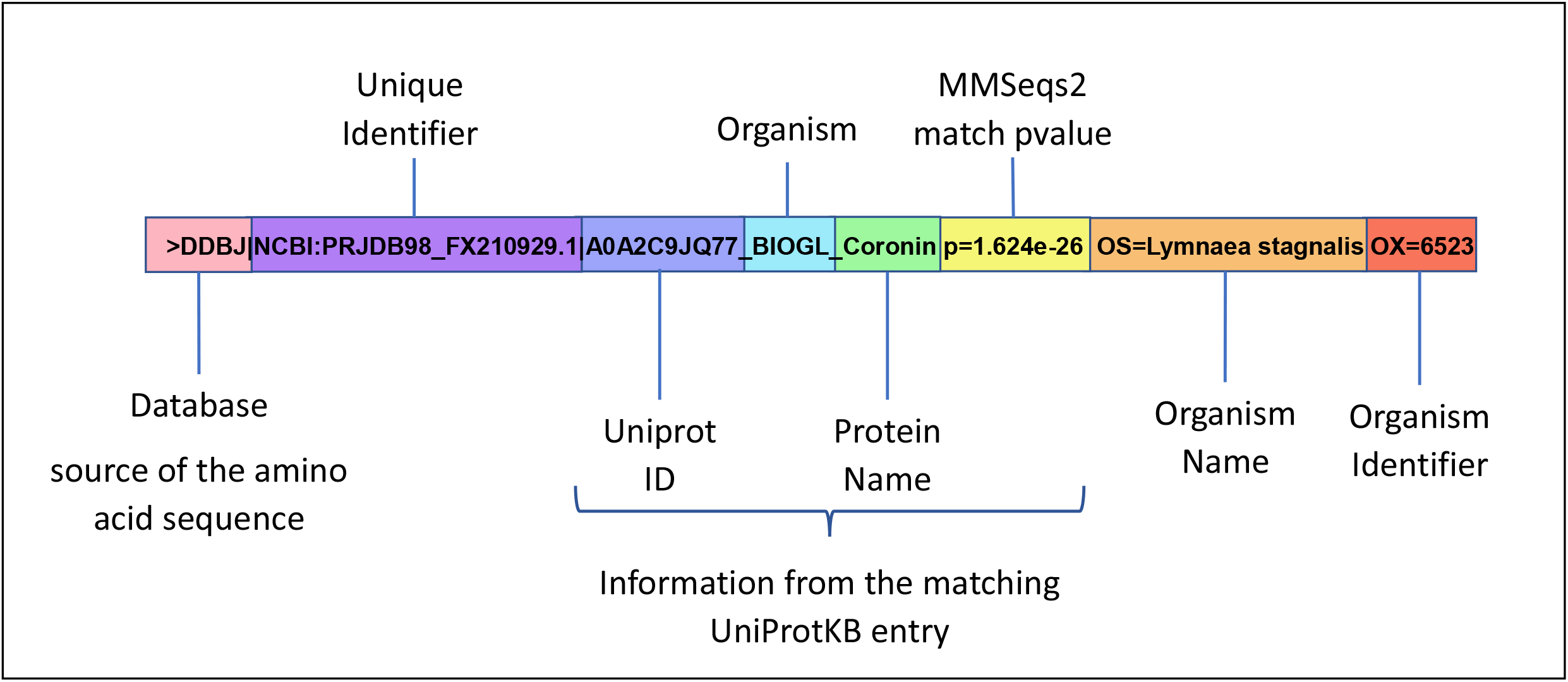
Design of the header composition used for the LymSt PDB following the UniProtKB parsing rules for FASTA headers.

To compare our database with the non-specific database, we have prepared a molluscan database with all proteins identified from our LymCNS-PDB database using the amino acid sequences of the matching molluscan species from UniProtKB. For an optimal comparison, both databases have the same size and contain the same number of proteins.

## Results

### Construction of the LymCNS-PDB

The NCBI PRJDB98 transcriptome dataset (Sadamoto et al., 2012) which contains 116,265 transcripts was analysed with a Trinity TransDecoder (v 5.5.0) to predict coding regions in the present assemblies. 22,180 ‘transdecoded’ transcripts were then used for homology searches with MMSeqs2 against 211,200 protein entries from the UniProtKB database with the preference for *Lymnaea stagnalis*>Euthyneura>Mollusca. We were able to match 16,142 transcripts to 9,628 proteins from the UniProtKB database (Figure 2).

**Figure 2.**
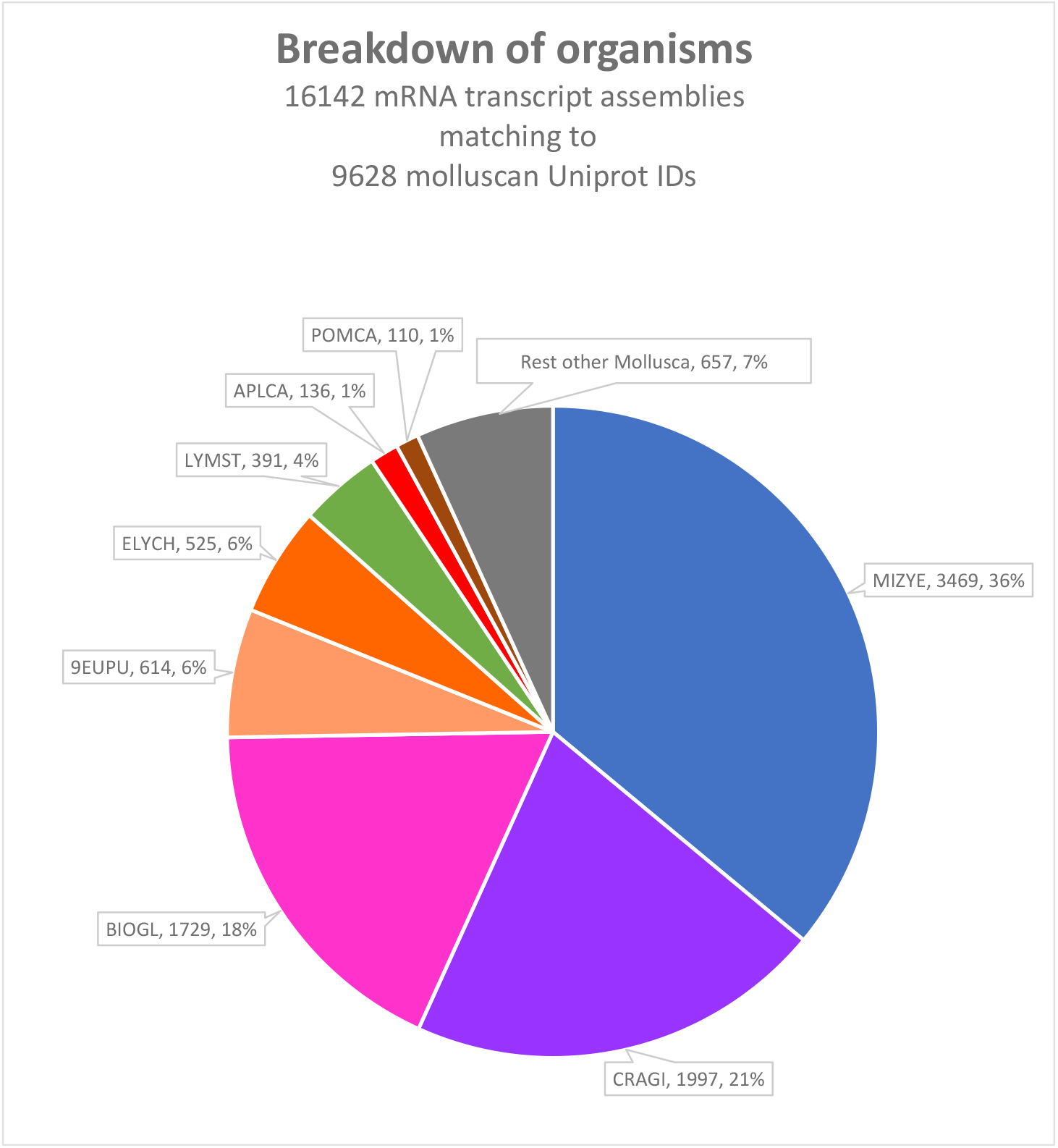
Breakdown of organisms showing the percentual distribution of matching transcript assemblies of NCBI PRJDB98 transcriptome dataset to proteins of different organisms within the UniProtKB Mollusca database. MIZYE (*Mizuhopecten yessoensis*); CRAGI (*Crassostrea gigas*); BIOGL (*Biomphalaria glabrata*); 9EUPU (*Eupulmonata*/ majority *Arion vulgaris* with 75%); ELYCH (*Elysia chlorotica*); LYMST (*Lymnaea stagnalis*), APLCA (*Aplysia californica*); POMCA (*Pomacea canaliculata*).

Three quarters of all matching entries were from organisms with a large number of available protein sequences in UniProtKB: *Mizuhopecten yessoensis* (MIZYE; *Yesso scallop* with 22,614 protein entries matching to 4781 PRJDB98 transcripts of 3,469 unique proteins; *Crassostrea gigas* (CRAGI; Pacific oyster) with 27,077 protein entries matching to 2,672 PRJDB98transcripts of 1997 unique proteins and *Biomphalaria glabrata* (BIOGL), a species of air-breathing freshwater snail, with 31,775 protein entries matching to 3,183 PRJDB98 transcripts of 1,729 unique proteins.

A rather low number of matching entries were from the following molluscan organisms: A mixture of several *Eupulmonata* (9EUPU), with the majority of protein entries (75%) from *Arion vulgaris* within this taxonomic clade of air-breathing snails, with 65368 protein entries matching to 1072 PRJDB98 transcripts of 614 unique proteins; *Elysia chlorotica* (ELYCH; eastern emerald elysia) with 23887 protein entries matching to 928 PRJDB98 transcripts of 525 unique proteins; *Lymnaea stagnalis* (LYMST; great pond snail) with 442 protein entries matching to 1245 PRJDB98 transcripts of 391 unique proteins; *Aplysia californica* (APLCA; california sea hare) with 443 protein entries matching to 755 PRJDB98 transcripts of 136 unique proteins, *Pomacea canaliculata* (POMCA; golden apple snail) with 21,514 protein entries matching to 144 PRJDB98 transcripts of 110 unique proteins, and the rest of 237,428 molluscan protein entries matching to 1,029 PRJDB98 transcripts of 657 unique proteins (Figure 2 and Table 1).

**Table 1.**
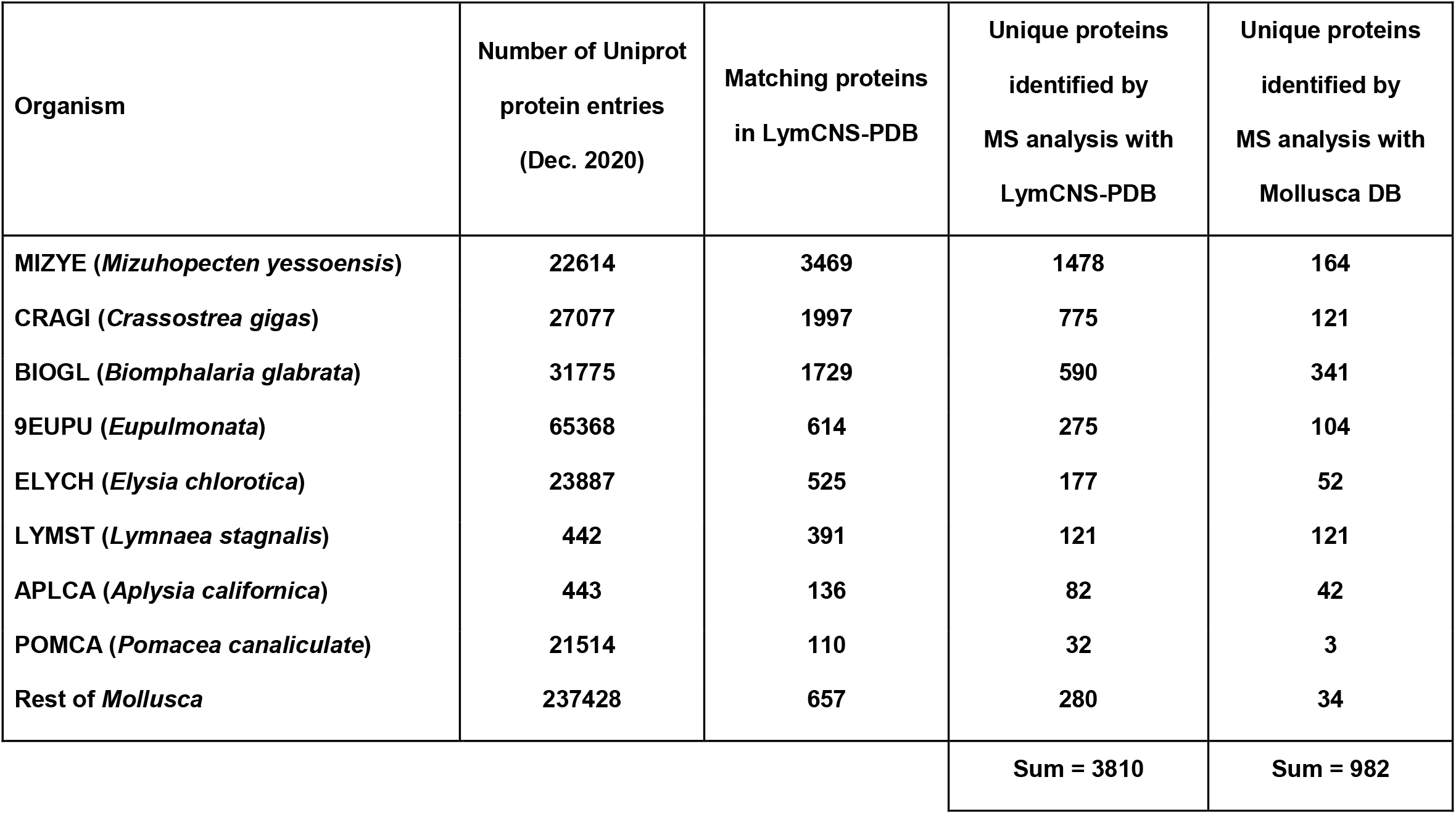
Overview of the number of protein entries for the respective organisms in UniProtKB Database, number of matching proteins in the resulting LymCNS-PDB database, number of unique proteins identified by MS analysis using the LymCNS-PDB database in comparison to the number of unique proteins identified by MS analysis using a molluscan database from UniProtKB.

| Organism | Number of Uniprot<br>protein entries<br>(Dec. 2020) | Matching proteins<br>in LymCNS-PDB | Unique proteins<br>identified by<br>MS analysis with<br>LymCNS-PDB | Unique proteins<br>identified by<br>MS analysis with<br>Mollusca DB |
| --- | --- | --- | --- | --- |
| MIZYE (*Mizuhopecten yessoensis*) | 22614 | 3469 | 1478 | 164 |
| CRAGI (*Crassostrea gigas*) | 27077 | 1997 | 775 | 121 |
| BIOGL (*Biomphalaria glabrata*) | 31775 | 1729 | 590 | 341 |
| 9EUPU (*Eupulmonata*) | 65368 | 614 | 275 | 104 |
| ELYCH (*Elysia chlorotica*) | 23887 | 525 | 177 | 52 |
| LYMST (*Lymnaea stagnalis*) | 442 | 391 | 121 | 121 |
| APLCA (*Aplysia californica*) | 443 | 136 | 82 | 42 |
| POMCA (*Pomacea canaliculate*) | 21514 | 110 | 32 | 3 |
| Rest of *Mollusca* | 237428 | 657 | 280 | 34 |
|  |  |  | Sum = 3810 | Sum = 982 |

### LC-MS based proteomics analysis

In order to demonstrate the suitability of our newly created protein database (LymCNS-DB), we performed an MS-based proteomics experiment of the CNS from *Lymnaea stagnalis*, which resulted in the identification of 3810 unique proteins. The identified proteins were derived from the following molluscan organisms with significant matching homology to the corresponding amino acid sequences of the *Lymnaea stagnalis* transcripts from the PRJDB98 dataset: *Mizuhopecten yessoensis* (MIZYE; Yesso scallop) with 1,478; *Crassostrea gigas* (CRAGI; Pacific oyster) with 775; *Biomphalaria glabrata* (BIOGL) with 590; *Eupulmonata* (9EUPU) with 275; *Elysia chlorotica* (ELYCH; eastern emerald elysia) with 177, *Lymnaea stagnalis* (LYMST; great pond snail) with 121; *Aplysia californica* (APLCA; california sea hare) with 121; *Pomacea canaliculata* (POMCA; golden apple snail) with 32 and other molluscan organisms with 280 unique proteins (Figure 3 and Table 1). In contrast, when the same experimental data was analysed against the non-specific molluscan database, only 920 unique proteins were identified with 44 proteins from *Mizuhopecten yessoensis*, 121 proteins from *Crassostrea gigas*, 341 proteins from *Biomphalaria glabrata*, 104 proteins from *Eupulmonata*, 52 proteins from *Elysia chlorotica*, 121 proteins from *Lymnaea stagnalis*. 41 proteins from *Aplysia californica*, 3 proteins from *Pomacea canaliculate* and 34 proteins from other molluscan species (Table 1).

**Figure 3.**
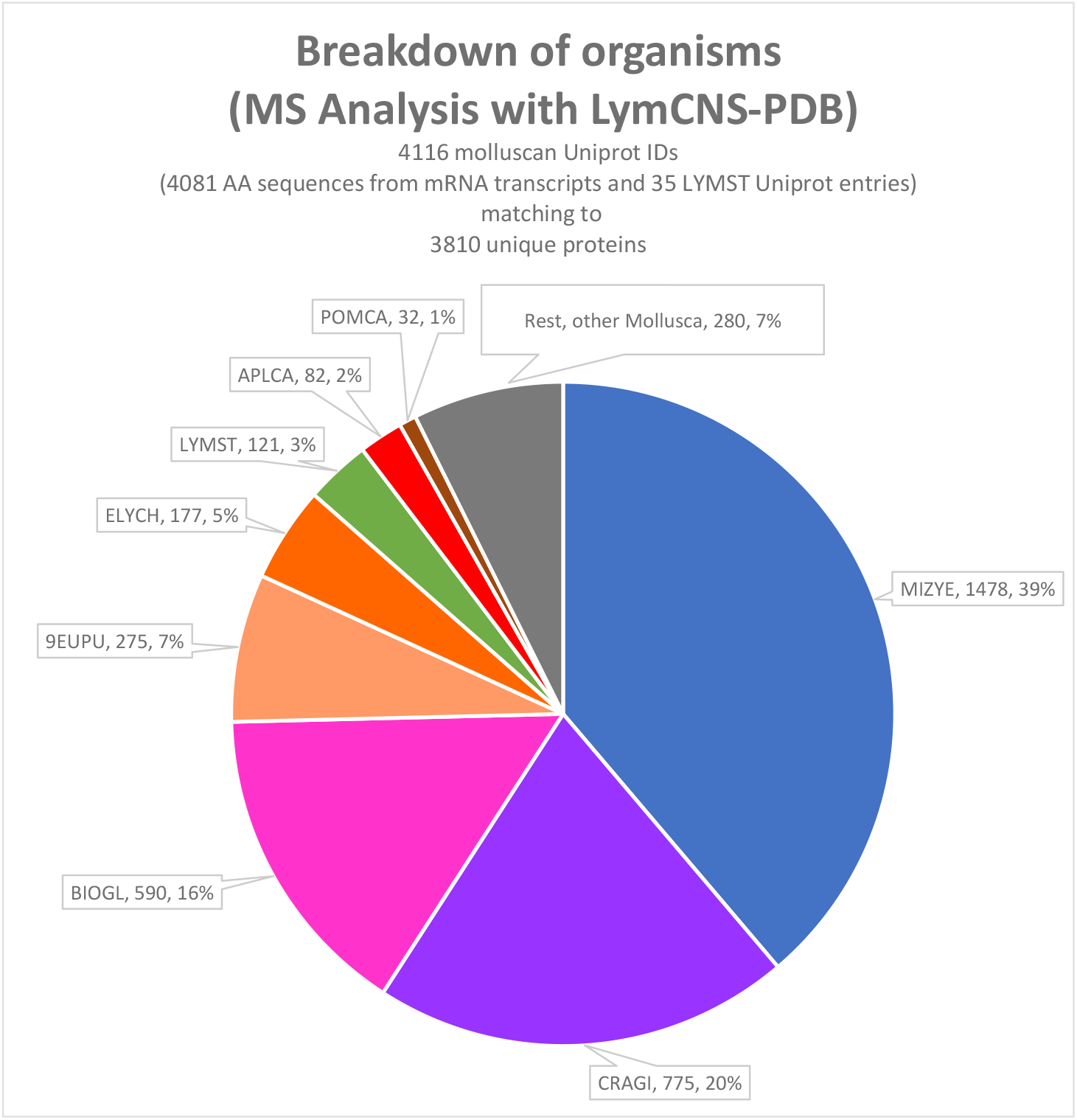
Percentual distribution of matching organisms of the identified proteins by LC-MS analysis using our *Lymnaea stagnalis* Proteomics database. MIZYE (*Mizuhopecten yessoensis*); CRAGI (*Crassostrea gigas*); BIOGL (*Biomphalaria glabrata*); 9EUPU (*Eupulmonata*/ majority *Arion vulgaris* with 75%); ELYCH (*Elysia chlorotica*); LYMST (*Lymnaea stagnalis*), APLCA (*Aplysia californica*); POMCA (*Pomacea canaliculata*).

## Discussion

The pond snail, *Lymnaea stagnalis*, is a highly successfully used invertebrate model organism in both basic and translational neuroscience research aimed at understanding the neural and circuit mechanisms of a variety of behaviours due to its numerically simple and well-characterized CNS (Benjamin 2008; Fodor et al. 2020; Rivi et al., 2020). However, the molecular characterization of the different functions of the CNS has been limited by a lack of a comprehensive proteomics database. In this investigation, we set out to develop an extensive proteomics database based on the mRNA transcript assemblies from the NCBI Bioproject PRJDB98. A recently published paper by Dong et al. 2021 presented a more comprehensive *L. stagnalis* transcriptomic database but included RNA transcripts from the ring without the buccal ganglia and the identified protein sequences were focused only on different ion channels (Dong et al., 2021). In our approach, we have developed a more general proteomics database that includes proteins involved in several CNS functions such as learning, fundamental decision making, motivational states, encoding hunger states and others. As previous published studies have identified the buccal ganglia as the location of circuitry involved in the expression of both appetitive and aversive memories (Ito et al., 2012; Marra et al., 2010), encoding hunger states (Crossley et al., 2016; Staras et al., 2003) as well as fundamental decision making (Crossley et al., 2018), we developed a proteomics database created from a transcriptomic database that included RNA transcripts from the whole CNS including the “learning” buccal ganglia and the ring. For this reason, we used the *L. stagnalis* transcriptomics database published by Sadamoto et al. 2012 as this database contained the transcriptome from the whole CNS including the “learning” buccal ganglia and the cerebral ring (Sadamoto et al., 2012).

By matching all mRNA transcript assemblies from the NCBI Bioproject PRJDB98 to all available molluscan proteins on the UniProtKB database, we succeeded in creating the proteomics database LymCNS-PDB with 9,628 Proteins, containing the translated amino acid sequences of their respective mRNAs from the *Lymnaea stagnalis* CNS, as well as obtaining all the other information (e.g., protein name) from their matching molluscan counterparts by the use of UniProtKB. Most of the matches to certain molluscan species were due to the large number of available protein sequences of this organism in UniProtKB. Species that have a smaller number of identified proteins have a greater match compared to those with a larger number of identified proteins. For example, *Mizuhopecten yessoensis* with 22,614, *Crassostrea gigas* with 27,077 and *Biomphalaria glabrata* with 31,775 protein entries in UniProtKB were matching to 3,469, 1,997 and 1,729 PRJDB98 sequences, respectively. These matches represent 75% of all the matching proteins (Figure 2), even if two of these three organisms, *Mizuhopecten yessoensis* and *Crassostrea gigas*, have the most distant phylogenetic relationship to *Lymnaea stagnalis* amongst all of the matching organisms (Figure 4). On the other hand, *Eupulmonata*, that has a very high number of 65,368 UniprotKB entries, only matches to 614 UniprotKB entries even though it has a closer phylogenetic relationship to *Lymnaea stagnalis*. Whereas, the group with a very high number of 65,368 UniprotKB entries, *Eupulmonata* (with approx.49,000 UniProtKB entries from *Arion vulgaris*) is only matching to 614 PRJDB98 sequences, with an even closer phylogenetic relationship to *Lymnaea stagnalis* (Figure 4).

**Figure 4.**
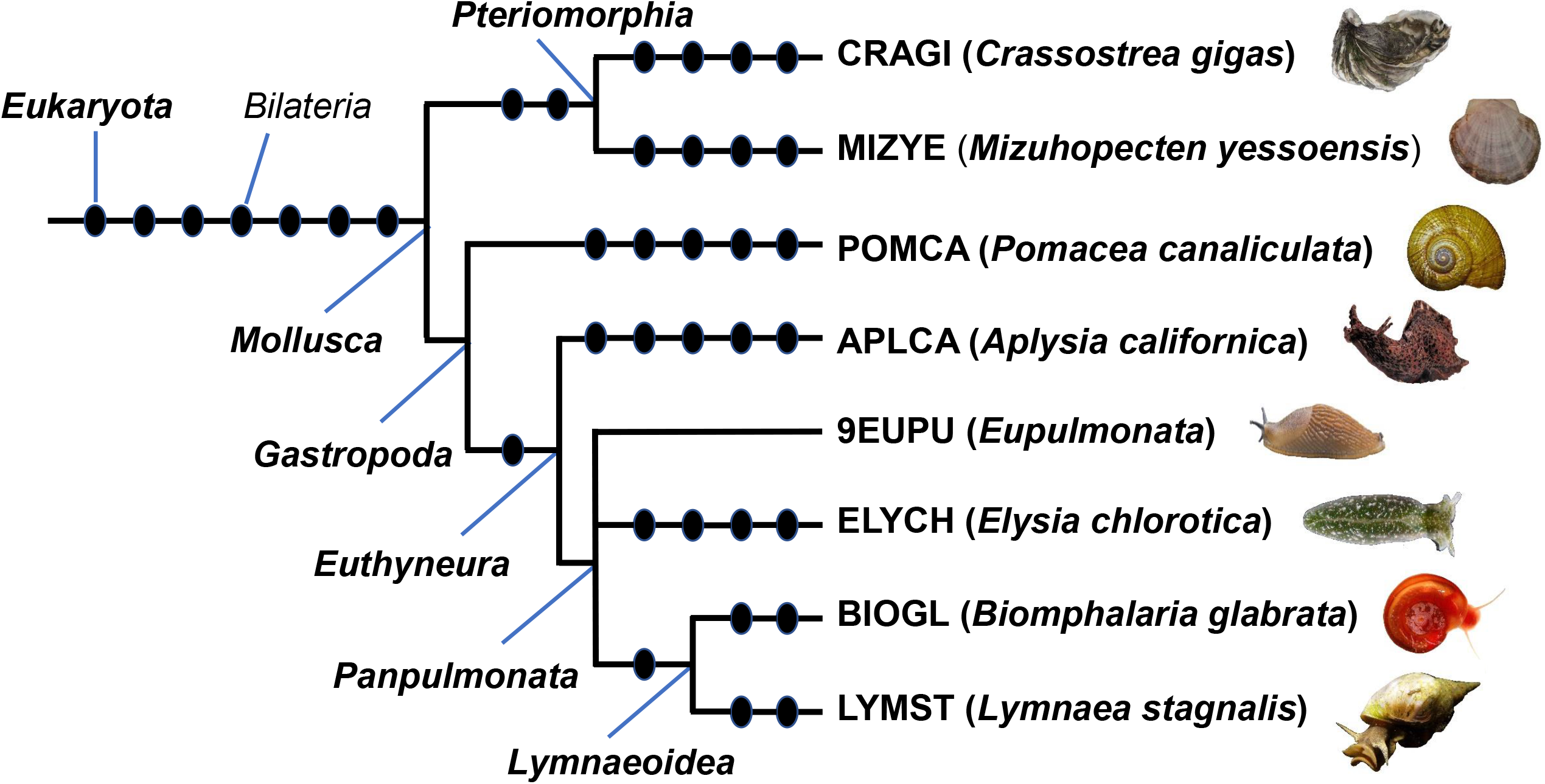
phyloT V2 (https://phylot.biobyte.de/) generated phylogenetic tree of the organisms with matching entries to the *Lymnaea stagnalis* transcripts of the NCBI PRJDB88 dataset. The Tree is generated, based on the NCBI taxonomy database, using MIZYE, CRAGI, BIOGL, 9EUPU, ELYCH, LYMST, APLCA and POMCA as NCBI tree elements. All pictures from https://commons.wikimedia.org

The number of protein identifications was increased by performing a pre-fractionation of the tryptic peptides from the CNS of *Lymnaea stagnalis* using immobilized pH gradient gels (ipg-gels) before analysis of the fractions by nanoLC-MS (Eravci et al., 2014).

Analysis of the MS/MS fragmentation spectra using the LymCNS-PDB led to an identification of 3,810 unique proteins that represents almost 40% of the entire LymCNS-PDB database. To our knowledge, this is the highest number of protein identifications in a proteomics experiment in *Lymnaea stagnalis*.

The breakdown of organisms of the MS analysis with LymCNS-PDB (Figure 3) shows the same distribution among the matching organisms as in the breakdown of organisms showing the number of matches from all molluscan entries to the NCBI Bioproject PRJDB98 assemblies. The similarity of both distributions (Figures 2 and 3) indicates that there is no bias in the comparison of the MS/MS fragmentation spectra to the amino acid sequences in the LymCNS-PDB since the overall distribution of the matching organisms is the same as the entire database.

To compare our database with specific amino acid sequences for *Lymnaea stagnalis* with the approach of other studies that were using a non-specific database for the identifications of proteins from this species (Giusti et al., 2013; Rosenegger et al., 2010; Silverman-Gavrila et al., 2011), we prepared a molluscan database with all proteins from our LymCNS-PDB, but with the amino acid sequences of their matching molluscan counterparts from UniProtKB. After analysis of the same experimental data, which were previously used for the analysis with the LymCNS-PDB, the comparison with the unspecific molluscan database led to an identification of only 920 unique proteins, which are presumably containing orthologous sequences in evolutionally conserved regions, homologues to the sequences of certain tryptic peptides from the *Lymnaea stagnalis* CNS.

This comparison clearly showed the benefits of using a proteomics database with specific amino acid sequences for the organism under investigation, as was the case of LymCNS-PDB for the identification of proteins from the *Lymnaea* CNS, which showed a 4 times higher number of identified proteins compared with the non-specific molluscan database.

We have presented the most extensive proteomics database to date, specifically for proteins from the CNS of *Lymnaea stagnalis*, the LymCNS-PDB. We have successfully used this database, as a proof of principle, to identify proteins in the isolated *Lymnaea* CNS. The LymCNS-PDB database provides a valuable tool to open new avenues for future research on proteomics to identify and quantify a plethora of proteins, which are involved in the molecular mechanisms of different neurobiological functions in the CNS of *Lymnaea stagnalis* including learning and memory formation, aging, feeding patterns, defensive responses and neuro-hormonal behavioural circuits involved in reproduction.

## Acknowledgements

This work was funded by the Biotechnology and Biology Research Council grant BBSRC/BB/P00766X/1 (I.K., G.K., P.R.B.). The authors would like to thank Dr. Paul Johnston from the Freie Universität Berlin for the analysis of the NCBI PRJDB98 transcriptome dataset with Trinity TransDecoder and providing the transcoded dataset. For mass spectrometry, we acknowledge the assistance of the Core Facility BioSupraMol supported by the Deutsche Forschungsgemeinschaft (DFG).

## Contributions

SW performed the MMSeqs2 homology searches and constructed the LymSy-PDB under the supervision of FP. Dissections of nervous tissue and sample preparation were performed by AA and MC. LC-MS based proteomics analysis was conducted by ME and BK. Manuscript was written by ME and edited by AA, BK, IK, PRB and GK.

## Availability of data

PRoteomics IDEntifications Database (PRIDE, https://www.ebi.ac.uk/pride/):

Project accession: PXD025591

Project DOI: 10.6019/PXD025591

